# Defining the neural correlates of spontaneous theory of mind (ToM): An fMRI mega-analytic investigation

**DOI:** 10.1101/560953

**Authors:** Sara Boccadoro, Emiel Cracco, Anna R. Hudson, Lara Bardi, Annabel D. Nijhof, Jan R. Wiersema, Marcel Brass, Sven C. Mueller

**Affiliations:** Department of Experimental Clinical and Health Psychology, Ghent University, Ghent, Belgium; Department of Experimental Psychology, Ghent University, Ghent, Belgium; Institute of Cognitive Neuroscience Marc Jeannerod, CNRS / UMR 5229, 67 Bd Pinel, 69500 Bron, France; Social, Genetic and Developmental Psychiatry centre, Institute of Psychiatry, Psychology and Neuroscience, King’s College London, UK

**Keywords:** Theory of Mind, fMRI, spontaneous mentalising, explicit mentalising, false belief, temporo-parietal junction

## Abstract

There is a major debate in the theory of mind (ToM) field, concerning whether spontaneous and explicit ToM are based on the same or two distinct cognitive systems. While there is extensive research on the neural correlates of explicit ToM, revealing consistent activation of the temporo-parietal junction (TPJ) and medial prefrontal cortex (mPFC), few studies investigated spontaneous ToM, with conflicting results, probably due to having small samples of participants. Here, we implemented a mega-analytic approach by pooling data from three fMRI studies, to achieve enough statistical power to better define the neurocognitive mechanisms underlying spontaneous ToM. Participants watched videos in which an agent acquires a true or false belief about the location of a ball. By analysing the blood-oxygen level dependent signal during the belief formation phase for false versus true beliefs, we found a cluster of activation in the right, and to a lesser extent, left posterior parietal cortex spanning the TPJ, but no mPFC activation. Region of interest (ROI) analysis on bilateral TPJ and mPFC confirmed these results and added evidence to the content-selectivity for false beliefs with positive content and asymmetry in laterality of the spontaneous ToM system. Interestingly, the whole brain analysis, supported by an overlap with brain maps, revealed maximum activation in areas involved in visuospatial working memory and attention switching functions, such as the supramarginal gyrus, the middle temporal gyrus, and the inferior frontal gyrus. Taken together, these findings suggest that spontaneous and explicit ToM rely on partially overlapping brain systems. However, spontaneous ToM tasks also show clear differences with explicit ToM tasks.

## 1. Introduction

Humans are a social species and so, by definition, regularly engage in social interactions, which require the ability to understand and predict the goals, beliefs, desires, thoughts and behaviours of other people. This fundamental ability is called Theory of Mind (ToM) and is crucial into representing others’ mental states. Traditionally, ToM has been investigated using false belief tasks, mainly the ‘Sally-Anne’ false belief task (Baron-Cohen, Leslie & Frith, 1985; Wimmer & Perner, 1983), which *explicitly* asks participants to reason about other people’s mental states. In the Sally-Anne paradigm, participants watch a scene in which a character, Sally, places an object in a box before leaving the scene. After Sally leaves, another character Anne moves the object to a different box. When Sally re-enters the scene, participants are asked to indicate in which box they think Sally will look for the object. Indicating the correct box requires the capacity to represent Sally’s false belief. This line of research suggests that ToM requires executive functions to suppress one’s own belief, which emerge later in development and, thus, only children aged four years old and above tend to pass the classic false belief task (Wellman, Cross & Watson, 2001; Wimmer & Perner, 1983). However, following research has challenged this idea, showing that infants before this age, much like adults, can already represent others’ belief in a *spontaneous* way, when they are not explicitly required to do so or when others’ mental states are irrelevant for their goals (Clements & Perner, 1994; Kovács, Téglás & Endress, 2010; Onishi & Baillargeon, 2005; Schneider, Slaughter & Dux, 2017; Southgate, Senju & Csibra, 2007).

This spontaneous ToM ability can be investigated with tasks in which participants are not specifically instructed to think about others’ mental states. For example, Kovács and colleagues (2010) developed a novel behavioural object detection task, where participants had to watch a video depicting an agent (a smurf) acquiring information about the location of an object (a ball) that could be behind an occluder or not. Here, the belief of the agent and that of the participant could either match, in the True Belief condition (the participant and the agent believe the object to be at the same location), or differ, in the False Belief condition (the participant and agent believe the object to be at different locations). Participants were never asked to consider the belief of the agent and were only asked to press a button when the ball was present after the occluder fell. Importantly, whether the ball is present or absent is completely random in this task. Nevertheless, participants were biased by the beliefs of the agent, resulting in faster responses when the agent believed the ball was present, even when they themselves knew the ball should not be present.

With the advent of spontaneous ToM tasks the crucial question arose whether explicit and spontaneous ToM rely on similar or different cognitive systems. The first position, supported by Carruthers (2016), mantains that there is a single ToM system, which sometimes is working spontaneously and sometimes is controlled to voluntarily anticipate and understand other people’s behaviour, with the involvement of executive functions. The latter account, postulated by Apperly and Butterfill (2009), suggests the existence of two distinct systems, one that is present early in life, is efficient and fast and supports spontaneous ToM, and a later-developing, slower, flexible and more cognitively demanding system that supports explicit ToM.

One potential way to tease apart these two positions might be to investigate the neural correlates of explicit and spontaneous ToM and test whether they are based on overlapping or distinct brain networks. MRI studies on explicit ToM reveal consistent involvement of a pattern of brain regions forming the so-called “ToM network”, including the temporo-parietal junction (TPJ), the superior temporal sulcus (STS), the precuneus (PC), the temporal poles and the medial prefrontal cortex (mPFC) (Fletcher et al., 1995; Gallagher et al., 2000; Ruby & Dacety, 2003; Saxe & Kanwisher, 2003). Furthermore, meta-analytic evidence of explicit ToM suggests consistent activation of the TPJ and the mPFC (Decety & Lamm, 2007; Molenberghs, Johnson, Henry & Mattingley, 2016; Schurz, Radua, Aichhorn, Richlan & Perner 2014; Van Overwalle, 2009). Neural data on spontaneous ToM tasks in much smaller samples of healthy participants, by contrast, has obtained mixed findings to date (Bardi, Desmet, Nijhof, Wiersema & Brass, 2017 (N=22); Kovács, Kühn, Gergely, Csibra & Brass 2014 (N=15); Naughtin et al., 2017 (N=22); Schneider, Slaughter, Becker & Dux, 2014 (N=16)). For example, following up on their earlier study (Kovács et al., 2010), Kovács and colleagues (2014) and Bardi and colleagues (2017) reported enhanced TPJ activation for false than for true beliefs in the belief formation phase (namely when participants form a belief about the location of the object). However, contrary to explicit ToM studies, no mPFC activation during the belief formation phase for false versus true belief emerged. On the other hand, Schneider and colleagues (2014) did not find activation of the TPJ in a spontaneous ToM task, but only recruitment of the posterior cingulate and STS. In addition, Naughtin and colleagues (2017) reported higher activity in a more extensive network comprising the right TPJ, precuneus, left MFG and right STS for false belief trials relative to no belief trials. Therefore, more confirmatory evidence regarding the spontaneous ToM network is needed, especially regarding the role of the TPJ and the mPFC. Of crucial importance, the limited number of participants recruited for spontaneous ToM studies is a great limitation of power in the investigation of the neural correlates of spontaneous ToM with fMRI, thus reducing the ability to find a consistent pattern of activation and limiting replication. In particular, the absence of the mPFC in most spontaneous ToM studies might be due to a lack of power in detecting its activity. Next to the general question about the neural networks supporting spontaneous ToM, there are a number of additional fMRI findings in this task that require replication with sufficient statistical power.

Kovács and colleagues (2014), for example, observed an asymmetry in false belief activation in positive versus negative content, showing that the right TPJ was only active when the agent has a false belief with positive content (i.e., the agent believes that the ball is behind the occluder). They concluded that spontaneous ToM in contrast to explicit ToM might be restricted to the representation of false beliefs about the presence but not the absence of an object. Both Bardi and colleagues (2017) and Nijhof and colleagues (2018) replicated this content-selectivity of the rTPJ. However, replication of these findings in a bigger sample is needed to support this sensitivity to content of false belief.

Lateralization and functional asymmetry of the TPJ is another important current debate in the ToM field, with some studies claiming that the right TPJ is more specifically involved in explicit ToM than its left counterpart (Aichhorn et al., 2009; Döhnel et al., 2012; Liu, Meltzoff & Wellman, 2009; Saxe, 2010) and others favouring a more bilateral activation of the TPJ in ToM tasks (Jenkins & Mitchell, 2010, Krall et al., 2015; Saxe & Kanwisher, 2003). More recently, Kovács and colleagues (2014), Bardi and colleagues (2017) and Nijhof, Bardi, Brass and Wiersema (2018) reported the recruitment of only the right TPJ in spontaneous ToM, but also left open the question about a possible role of the left TPJ, given the association between damage to the left TPJ and explicit ToM deficits, reported in lesion studies (Apperly, Samson, Chiavarino & Humphreys, 2004; Samson, Apperly, Chiavarino & Humphreys, 2004). In other words, previous studies have not directly tested the hypothesis that spontaneous ToM is right (versus left) lateralised.

Given the scarcity of studies investigating the spontaneous ToM system, contradictory findings, and the limited number of participants in each experiment, the present study sought to investigate the above hypotheses using a mega-analytic design in a sample of healthy participants. To this end, we combined data from three previous MRI studies that used the same spontaneous ToM task (in the same MRI scanner) to investigate the neural correlates of spontaneous false belief processing in a bigger sample (Bardi et al., 2017; Hudson, Van Hamme, Maeyens, Brass & Mueller [preprint, bioRxiv]; Nijhof et al., 2018). Using such an approach should result in a sufficiently large sample to overcome the power issue limiting previous findings. Indeed, the goal of this study was to resolve the three main issues that surround brain imaging work on spontaneous ToM, namely the potential involvement of the mPFC in the formation of belief, the content specificity of spontaneous ToM, and the lateralisation of TPJ activation. We had four predictions: firstly, we expected overlap with the brain regions activated during explicit ToM in the TPJ and mPFC. Indeed, if the absence of mPFC activity in False > True Belief during the belief formation phase in most spontaneous ToM studies is only due to a power issue, we would expect its activation in a larger sample. To look for overlap with explicit ToM, we performed region-of-interest (ROI) analyses on TPJ and mPFC. Secondly, we expected to replicate the asymmetry of TPJ activation in response to positive content, with higher TPJ activity in the False Belief with positive content condition. Thirdly, directly testing the issue of laterality, we predicted greater activity in the right TPJ than in the left TPJ, which would confirm the right (versus left) dominance for spontaneous ToM processing. Fourthly, to assess the consistency of our findings with the ToM literature, we compared our activation map to the average ToM map using Neurosynth (http://www.neurosynth.org/). Lastly, we wanted to explore the correlation between brain activity in the belief formation phase and behaviour in the outcome phase. We also included an exploratory psychophysiological interaction (PPI) analysis of TPJ connectivity.

## 2. Materials and methods

### 2.1 Participants

This mega-analysis includes samples from three different, independent studies conducted at Ghent University (Bardi et al., 2017, Hudson et al., [preprint, bioRxiv]; Nijhof et al., 2018), all of which used the same identical spontaneous ToM task but with different participants. In total, data from 74 healthy participants (19 males; mean age = 31.08 years, SD = 10.29) were available. Six subjects had to be excluded due to excessive movement (> 3 mm or 3° on any dimension), resulting in a total sample of 68 participants (17 males; mean age = 31.13 years, SD = 10.49). All participants had normal or corrected-to-normal vision, did not have any reported history of neurological disorders and gave written informed consent prior to the study. Handedness information was not available for all participants included in this study. All three included studies were approved by the Medical Ethics Review Board of Ghent University Hospital.

### 2.2 Task and stimuli

In all three included studies, participants performed a spontaneous ToM task, called the “Buzz Lightyear” task (Bardi et al., 2017; Nijhof et al., 2018), which is an adaptation of the task originally developed by Kovács et al. (2010). Participants laid in the MRI scanner while watching short videos and detecting an object at the end of each video. All movies consisted of two phases: the belief formation phase and the outcome phase. Each movie lasted 13.8 s. All movies started with the belief formation phase, in which an agent (*Buzz Lightyear*) placed a ball on a table in front of an occluder. The ball rolled behind the occluder at 3 s and after this, the movie could continue in four possible ways:

1. In the True Belief-Positive Content condition, the ball rolled out of the scene from behind the occluder and then rolled back behind it at 10 s in the presence of the agent, who then left the scene at 11 s. As a consequence, both the participant (P) and the agent (A) believed that the ball was behind the occluder (P+A+).
2. In the True Belief-Negative Content condition, after emerging from behind the occluder without leaving the scene, the ball rolled back behind the occluder and then left the scene at 10 s in the presence of the agent, who left the scene at 11 s. Therefore, both the participant and the agent believed that the ball was not behind the occluder (P-A-).
3. In the False Belief-Positive Content condition, the ball was behind the occluder when the agent left the scene at 6 s. Then the ball emerged from behind the occluder without leaving the scene, rolled back behind the occluder and finally left the scene at 11 s, when the agent was absent. Thus, the participant believed that the ball was not behind the occluder, whereas the agent wrongly believed that the ball was behind the occluder (P-A+).
4. In the False Belief-Negative Content condition, the ball rolled out of the scene while the agent was present. The agent left the scene at 9 s and, in his absence, the ball rolled back behind the occluder at 11 s. Therefore, the participant believed that the ball was behind the occluder, whereas the agent wrongly believed that the ball was not behind the occluder (P+A-).

In order to ensure that attention was maintained throughout the presentation of the movies, participants had to press a button with their left index finger as quickly as possible when Buzz left the scene.

In the outcome phase, at the end of each movie, the agent re-entered the scene, the occluder fell down and there could be two possible and equally probable outcomes, in which the ball could be either present or absent behind the occluder. At this point, participants had to press a button with their right index finger as quickly as possible, but only if the ball was present after the occluder fell. The presence (B+) or absence (B-) of the ball was completely independent of the belief formation phase, since the ball was present randomly in half of the trials. Thus, the ball could be expected or unexpected for both the participant and the agent. The combination of belief formation phase (P-A-; P+A+; P+A-; P-A+) and outcome phase (B+ and B-) resulted in eight different movies. Each movie was repeated 10 times, thus the task consisted of 80 trials presented in a randomised order in two blocks (fMRI runs) of 40 trials, with a short break in between, except for Hudson’s study, in which the 8 movies were repeated 8 times, thus resulting in 64 trials. The inter-trial interval was determined using a pseudo-logarithmic jitter with steps of 600 ms: half of the intervals were short (range from 200 to 2000 ms), one-third was intermediate (range from 2600 to 4400 ms) and one-sixth was long (range from 5000 to 6800 ms), with a mean inter-trial interval of 2700 ms. No instruction to reason about the agent’s belief was given to participants. Two studies (Bardi et al., 2017; Nijhof et al., 2018) included also an explicit ToM version of the task, always performed *after* the two spontaneous runs. Only the two spontaneous ToM runs were included in the present study.

### 2.3 fMRI data acquisition

Structural T1-weighted MRI images were acquired using a 3T Siemens Magnetom TrioTim MRI scanner. More specifically, 176 volumes of a T1-weighted MPRAGE high resolution structural image were acquired (repetition time (TR) = 2250 ms, echo time (TE) = 4.18 ms, image matrix = 256 × 256, field of view (FOV) = 256 mm, flip angle = 9°, slice thickness = 1 mm, voxel size = 1.0 × 1.0 × 1.0). The whole-brain T2*-weighted Echo Planar Images (EPI) sequence was identical across the three studies (TR = 2000 ms, TE = 28 ms, image matrix = 64 × 64, FOV = 224 mm, flip angle = 80°, slice thickness = 3.0 mm, voxel size = 3.5 × 3.5 × 3.0 mm, number of slices = 34).

### 2.4 fMRI data pre-processing

For the sake of consistency across studies, all data were pre-processed again with SPM8 software (Wellcome Department of Cognitive Neurology, London, UK) in MatLab (The Mathworks). The first four volumes for each EPI series were removed to allow magnetisation to reach a dynamic equilibrium. The pre-processing steps for the remaining volumes started with spatial realignment of the functional images using a rigid body transformation and then slice time correction of the realigned images with respect to the middle slice. The structural image of each subject was co-registered with the mean of the slice-time corrected images. During segmentation, the structural scans were brought in line with SPM8 tissue probability maps. The parameters estimated during the segmentation step were then used to normalise the functional images to standard MNI space. Lastly, the normalised functional images were resampled into 3 × 3 × 3 mm voxels and spatially smoothed with a Gaussian kernel of 8 mm (full-width at half maximum).

### 2.5 Behavioural data analysis

Reaction times (RT) were recorded for detection of the ball at the end of each movie. Behavioural data were analysed using IBM SPSS Statistics 25 (IBM Corp. Released 2017. IBM SPSS Statistics for Windows, Version 25.0. Armonk, NY, USA). For two participants, these data are missing due to technical problems during the recording. Therefore, data for these two participants were excluded from behavioural analyses. The behavioural analysis focused on the difference in RTs between the P-A- and the P-A+ condition, known as the *ToM Index* (Deschrijver, Bardi, Wiersema & Brass, 2016). This ToM Index is a measure of the influence of the agent’s belief on ball detection, so that a larger ToM Index indicates a larger influence. The reason why we did not include the P+ conditions in the behavioural analysis is that the effect of the agent’s beliefs on RTs in this task is known to be restricted to situations in which the participant believes the ball is not present (Deschrijver et al., 2016; Kovács et al., 2010).

### 2.6 fMRI data analysis

#### 2.6.1 Whole-brain analysis

First- and second-level analysis were carried out using SPM8. The subject-level statistical analyses were performed with a general linear model (GLM). Per run, the model contained four separate regressors for all combinations of Belief (true and false belief) and Belief Content (positive and negative content) in the belief formation phase, with durations of 9s modelled from the moment when the ball starts rolling on the table (3 s) to the moment when the agent re-enters the scene (12 s). In addition, eight regressors were added to model the outcome phase, (all possible combinations of Belief, Belief Content and Outcome), with duration of 0 s, modelled at the point when the occluder has completely fallen down and the presence or absence of the ball is revealed. In total, there were 12 regressors of interest per run. Furthermore, six subject-specific movement regressors calculated during the realignment step of preprocessing were added per run to account for head motion. All regressors were convolved with the canonical HRF.

The second-level analysis was conducted using a flexible factorial model with a between-subjects factor for experiment (to account for the fact that the data came from three different studies) and a within-subjects factor for condition (P+A+, P-A-, P-A+, P+A-). In order to identify regions involved in false belief tracking (belief formation phase), we computed our main contrast of interest (PxA, interaction contrast) as follows: False Belief (P- A+ and P+A-) > True Belief (P-A- and P+A+) and the reverse contrast for regions involved in true belief tracking. Additionally, we also computed the ToM Index (i.e., P-A+ > P-A-). Results for these whole-brain analyses were corrected for multiple comparisons using a p < 0.01 FWE whole-brain corrected threshold and minimal extent of 20 contiguous voxels. The reported coordinates correspond to the MNI coordinate system. Since our interest was to investigate the neural basis of false belief reasoning, we focused on the belief formation phase, in which the participant forms their own belief and that of the agent. Thus, the outcome phase was not taken into consideration for the analysis and no results are displayed for this phase.

To control for the influence of gender, an additional second-level analysis was conducted with this factor as a covariate of no interest. Finally, a last second-level analysis was conducted with the behavioural ToM Index as covariate to test the extent to which the neural ToM Index could be predicted by the behavioural ToM Index.

#### 2.6.2 ROI analysis

In addition to the whole-brain analysis, and to compare the findings from the spontaneous task with those coordinates commonly found in explicit ToM tasks, we performed a ROI-based analysis with *a priori* defined regions of interest (ROI), corresponding to the right and left TPJ and the mPFC, based on the meta-analysis by Kovács et al. (2014) (N=26 studies). Spheres with 6 mm radii centred on the right TPJ [56 −47 33], left TPJ [−56 −47 33] and mPFC [2 53 13] coordinates were created. Mean *β*s were extracted using the MARSBAR toolbox for SPM (Brett, Anton, Valabregue & Poline 2002). The obtained *β* values were entered into a repeated-measure ANOVA containing Participant (P- or P+) and Agent (A- or A+) belief as factors. Furthermore, we also included a Bayes Factor (BF) analysis with default JASP priors to calculate the likelihood of the data under the alternative hypothesis (False Belief > True Belief) relative to the null hypothesis (False Belief = True Belief). For example, a BF of 3 means that the data are three times more likely under the alternative hypothesis than under the null hypothesis, while a BF of 0.33 means the opposite (Rouder, Speckman, Sun, Morey & Iverson, 2009).

#### 2.6.3 Lateralisation

We carried out a second repeated-measure ANOVA to explore the degree to which TPJ activation was lateralised, by entering the *β* values of left and right TPJ with Participant, Agent and Location (left and right) as factors, and a third repeated-measure ANOVA to test the interaction between location and content of false belief, by entering the *β* values of left and right TPJ with Location (left and right) and Content of false belief (positive, P-A+ and negative P+A-) as factors.

#### 2.6.4 Cognitive decoding of brain activity

To estimate the consistency of our results with the average ToM activation found in the literature, the activation maps of both the False > True and True > False Belief PxA contrasts were entered in Neurovault (https://neurovault.org/collections/VQVUVUMR/; Gorgolewski et al., 2015). Neurovault allows us to use the cognitive decoding feature to compare the uploaded maps with the activation maps associated with various cognitive functions across many papers, using spatial correlations calculated in Neurosynth (http://neurosynth.org/). As output, this analysis reveals the cognitive functions whose activation maps are most correlated with the uploaded maps.

#### 2.6.5 Brain-Behaviour Relation

In addition to adding the behavioural ToM Index as a covariate, we also calculated the correlation of the neural ToM Index in the right and left TPJ ROIs with the behavioural ToM Index. The neural ToM Index was calculated by subtracting the β values for the P-A- condition to the β values for the P-A+ condition for each ROI.

#### 2.6.6 Functional connectivity (psychophysiological interaction, PPI)

To explore the effective connectivity between the right TPJ, a region we expected to be involved in belief reasoning, and the rest of the brain during the ToM task, a generalised psychophysiological interaction (PPI) analysis (McLaren, Ries, Xu & Johnson, 2012) was conducted. For this analysis, the same *a priori* TPJ coordinates as reported above together with the main activation peaks corresponding to the right TPJ (if any) of the False >True Belief contrast were used as source regions. A sphere of 6 mm radius was centred around each source region. The BOLD signal time series from each participant’s ROI were extracted and a voxel-wise PPI analysis was carried out. A second-level analysis was conducted in the same way as described for the task-based fMRI analysis. Results were corrected for multiple comparisons using a *p* < 0.01 FWE whole-brain corrected threshold and minimal extent of 20 contiguous voxels. All clusters reported (MNI coordinates) exceeded this cluster-corrected threshold. Results and discussion can be found in the Supplementary material.

## 3. Results

### 3.1 Behavioural results

Analysis of the ToM Index with a paired t-test revealed a significant difference between the P-A+ and the P-A- conditions, (*t*(66) = −4.57, *p* < 0.01) confirming the biasing effect of the agent’s belief on participants’ responses, so that participants responded faster when the agent believed the ball was present than when the agent believed the ball was absent (Figure 1).

**Figure 1.**
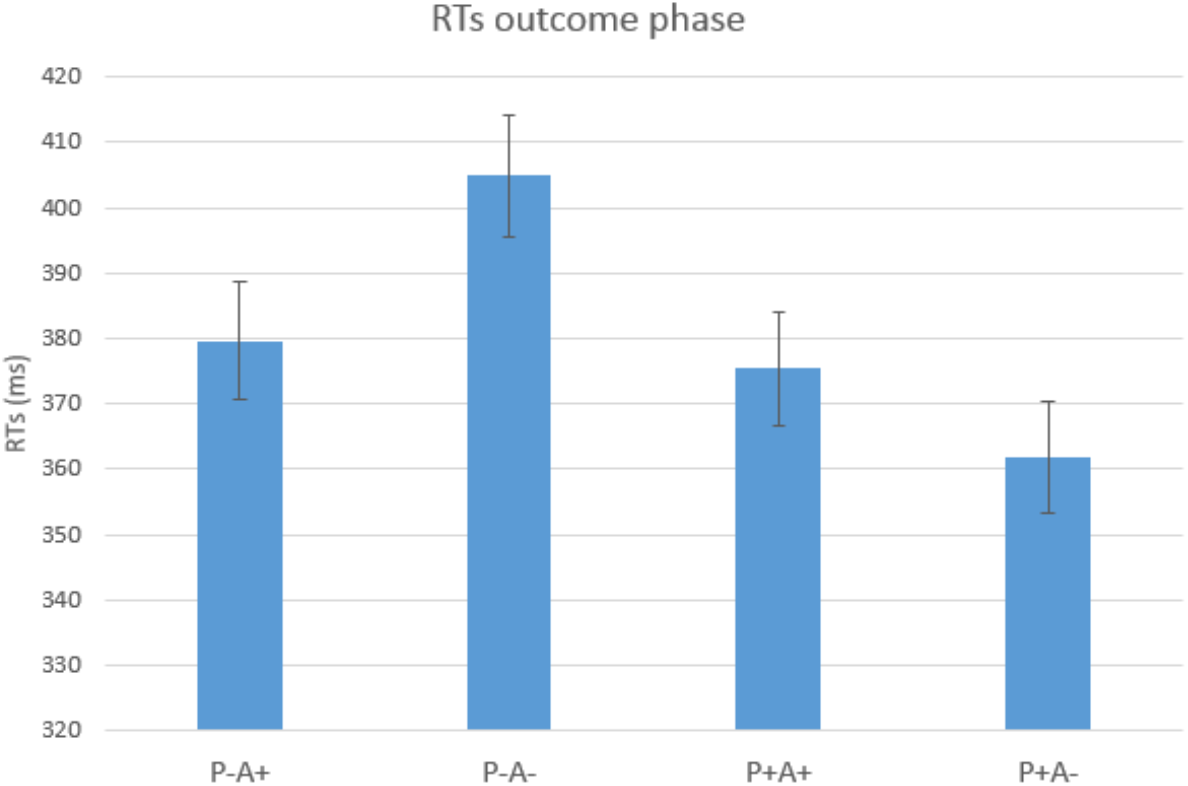
Behavioural analysis. The graphs show the reaction times (RTs) of the outcome phase for all the four different conditions (P-A+, P-A-, P+A+ and P+A-). The P-A+ and P+A- conditions are False Belief conditions and the P-A- and P+A+ are True Belief conditions. Error bars represent ± 1 standard error.

### 3.2 fMRI results

#### 3.2.1 Whole brain analysis

Our first aim was to identify the brain regions involved in spontaneous ToM. The PxA False > True Belief contrast yielded activation in a large cluster spanning the temporo-parietal cortex including the TPJ. Closer inspection of this cluster revealed that two of the three strongest activation peaks were located in the temporo-parietal cortex, with one peak in the middle temporal gyrus (MTG) (MNI coordinates [xyz]: 60 −52 1) and another in the supramarginal gyrus (SMG) (coordinates: 54 −37 46), while a third peak (coordinates: 12 −73 4) belonged to a more occipital region (the lingual gyrus). In addition to this temporo-parietal cluster, the PxA False > True Belief contrast also revealed activity in the right inferior frontal gyrus (IFG), the left inferior parietal lobule (IPL), the left middle temporal gyrus (MTG) and the right middle frontal gyrus (MFG). The same pattern of activity was found for the ToM Index F > T Belief contrast, except that the lingual gyrus peak found in the PxA analysis was now replaced by a peak in the angular gyrus (AG) (coordinates: 39 −46 37) (Table 1, Figure 2, Figure 3).

**Table 1.**
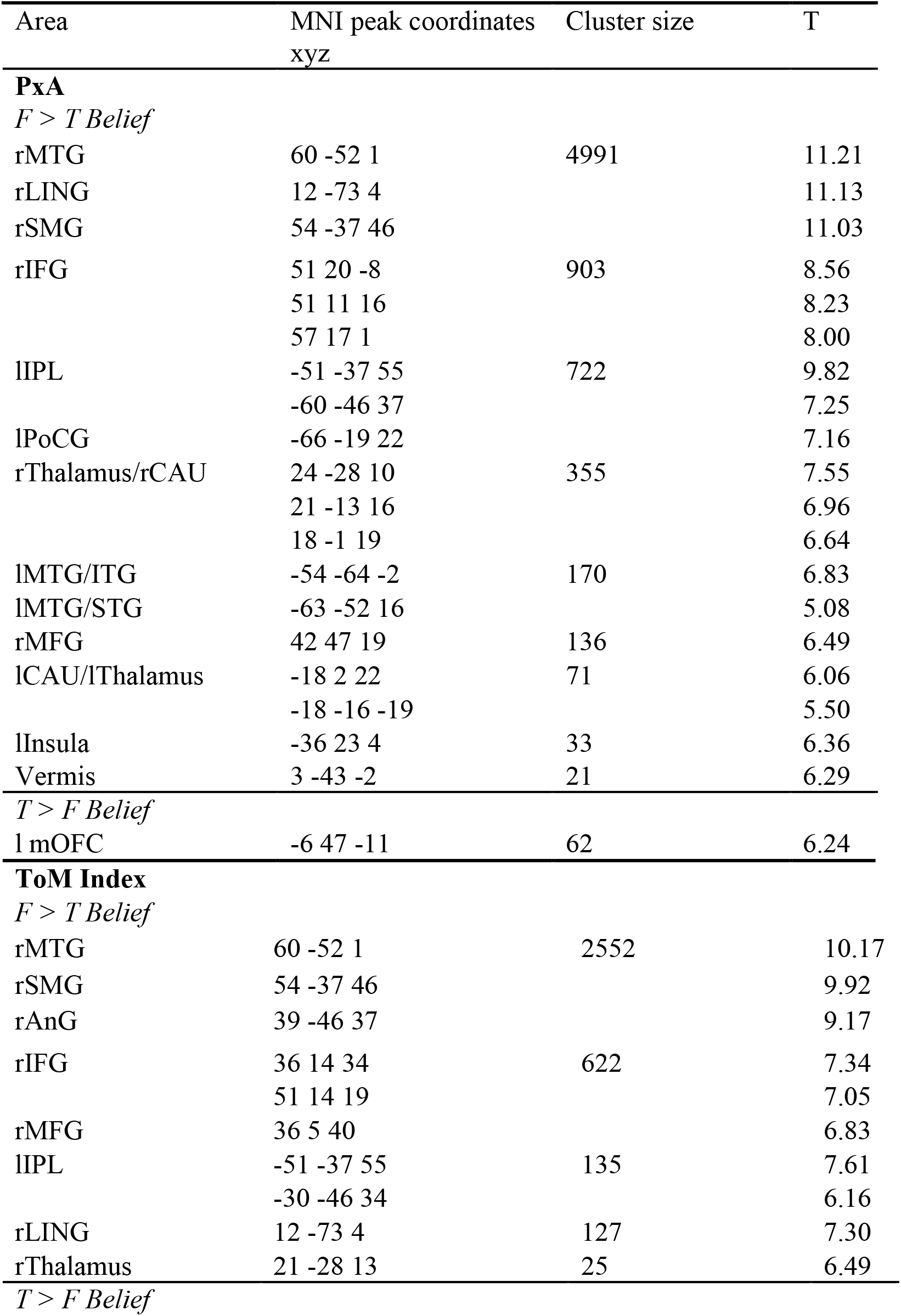
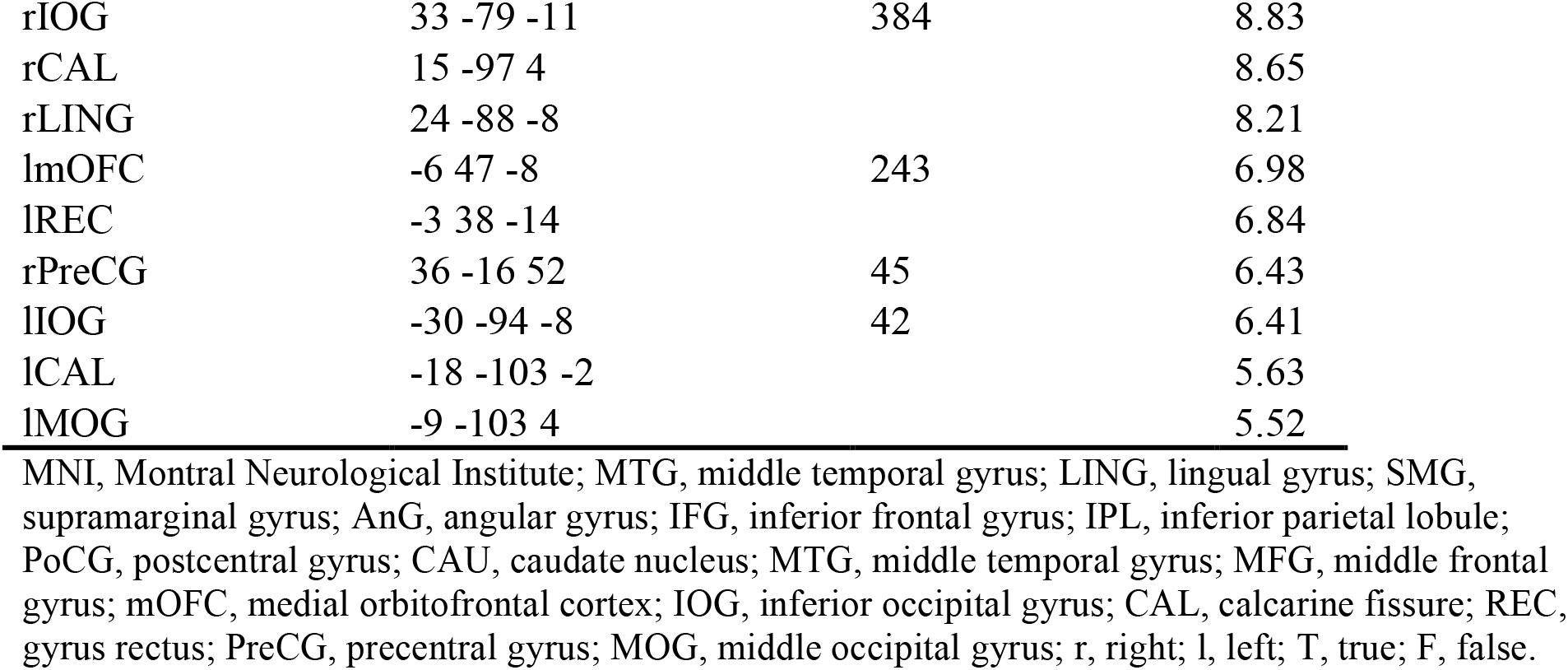
Peaks of activation in the belief formation phase.

**Figure 2.**
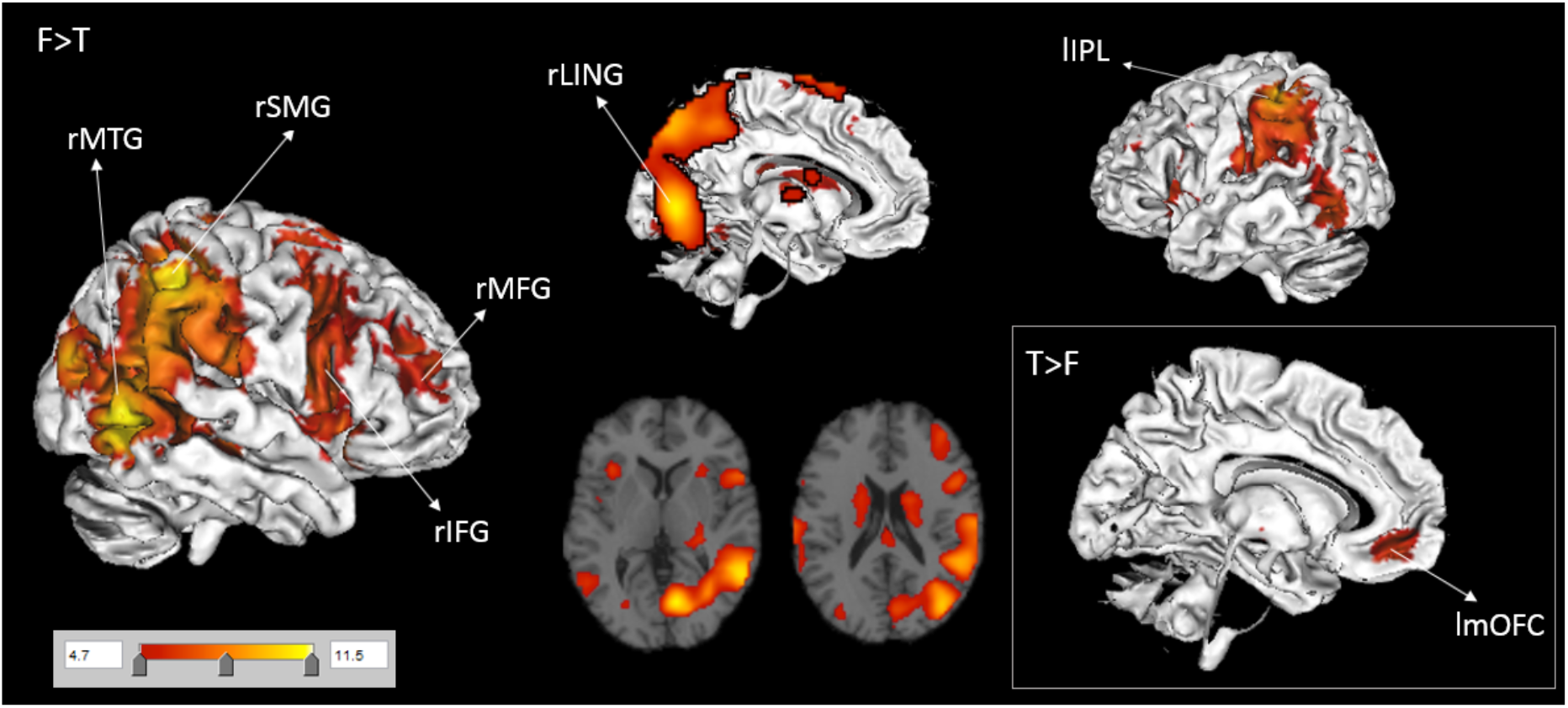
Results of whole-brain analysis for the PxA analysis. Figures were visualised with MANGO software (http://ric.uthscsa.edu/mango/). The figure displays the main areas activated during the spontaneous ToM task by the False Belief > True Belief condition (F > T) and the True > False Belief (T>F; bottom right panel) for the PxA contrast. The regions activated are coloured in red and yellow. SMG, supramarginal gyrus; MTG, middle temporal gyrus; LING, lingual gyrus; IFG, inferior frontal gyrus; MFG, middle frontal gyrus; IPL, inferior parietal lobule, mOFC, medial orbitofrontal cortex; r, right; l, left.

**Figure 3.**
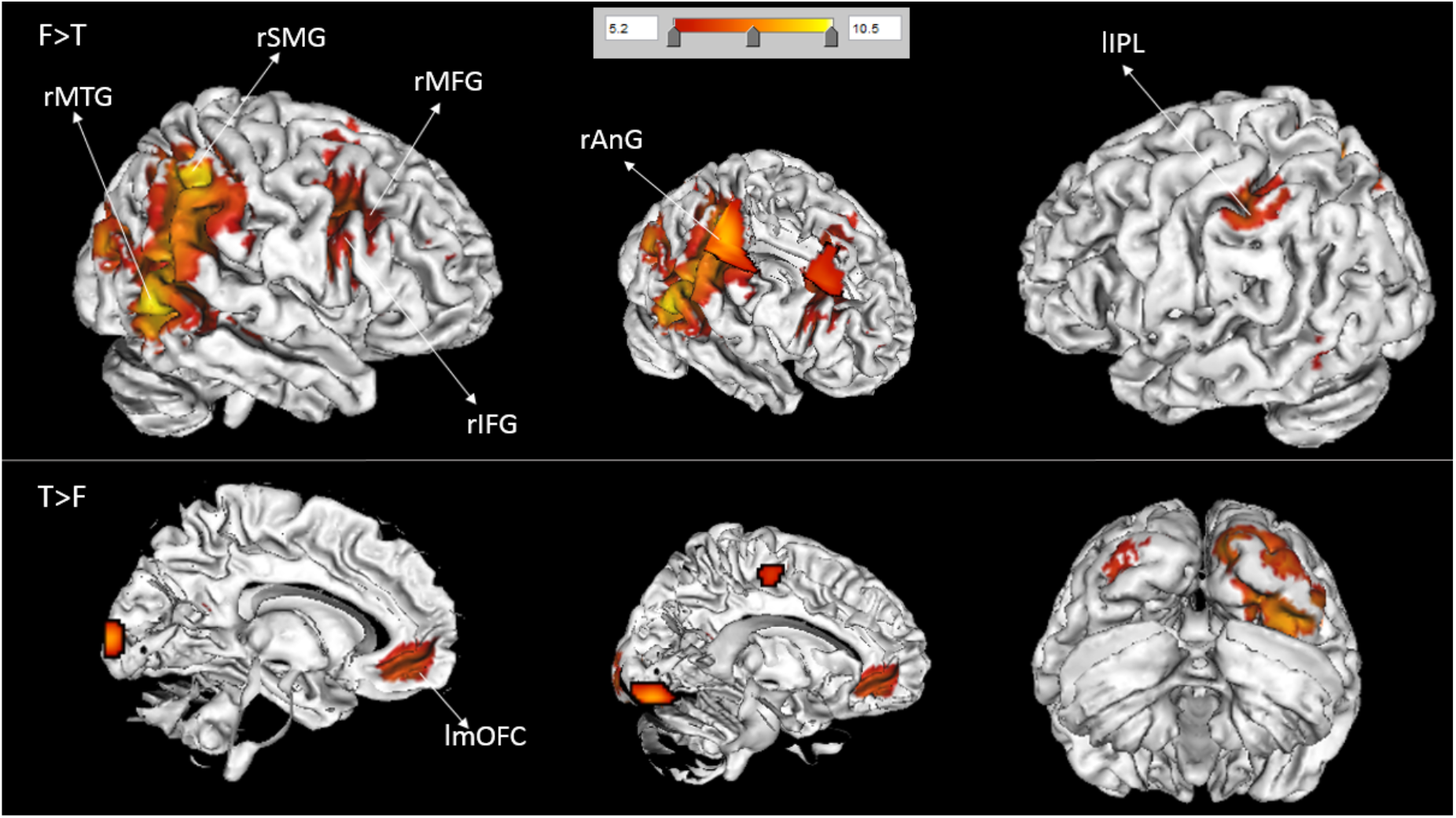
Results of whole-brain analysis for the ToM Index analysis. The figure displays the main areas activated during the spontaneous ToM task by the False Belief > True Belief condition (top panel) and True > False Belief condition (bottom panel) for the ToM Index contrast. The regions activated are coloured in red and yellow. SMG, supramarginal gyrus; MTG, middle temporal gyrus; IFG, inferior frontal gyrus; MFG, middle frontal gyrus; AnG, angular gyrus; IPL, inferior parietal lobule; mOFC, medial orbitofrontal cortex; r, right; l, left.

#### 3.2.2 TPJ and mPFC ROI analysis

To examine consistency with the broader literature, compare findings for the spontaneous ToM system with the explicit ToM system and verify if we can replicate the asymmetry in response to positive versus negative content, we also explored two a priori ROIs previously reported to be involved in belief processing, namely the bilateral TPJ and the mPFC. The results are displayed in Figure 4. A main effect of agent’s belief for the rTPJ, (*F*(1,67) = 8.04, *p* = 0.006) emerged. The rTPJ was more strongly activated when the agent believed the ball was present (A+) than when he believed the ball was absent (A-). This also resulted in a significant interaction between agent and participant for both TPJ regions (right: *F*(1,67) = 47.50, *p* < 0.001; left: *F*(1,67) = 28.95, *p* < 0.001), indicating stronger activation in bilateral TPJ in the False Belief conditions (P-A+ and P+A-) relative to the True Belief conditions (P-A- and P+A+).

**Figure 4.**
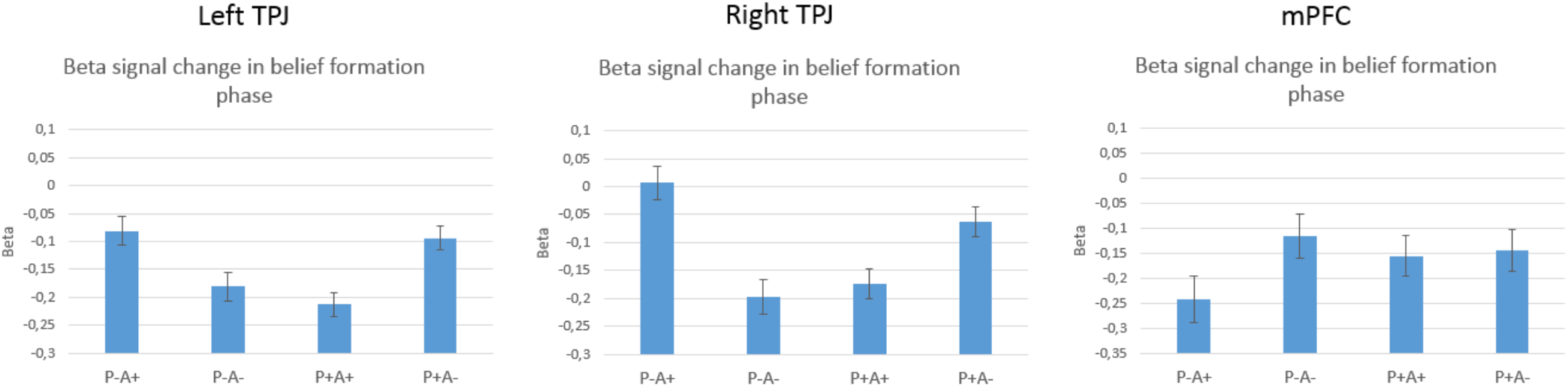
ROI analysis. The graphs show the beta signal change in the belief formation phase for all the four different conditions (P-A+, P-A-, P+A+ and P+A-) in the left TPJ, right TPJ and mPFC ROIs respectively. The P-A+ and P+A- conditions are False Belief conditions and the P-A- and P+A+ are True Belief conditions. Error bars represent ± 1 standard error.

A main effect of agent’s belief also emerged for the mPFC (*F*(1,67) = 9.52, *p* = 0.003) in the opposite direction to the TPJ. Indeed, the mPFC was more activated when the agent believed the ball was absent (A-) than when he believed the ball was present (A+). No significant interaction effect emerged for the mPFC (*F*(1,67) = 2.86, *p* = 0.097).

Bayesian analysis revealed a BF of 9.50 × 10^6 for the rTPJ and of 30,005 for the lTPJ, indicating strong evidence in favour of stronger activity in the right and left TPJ in the False Belief condition than in the True Belief condition. By contrast, for the mPFC the BF was 0.052, indicating strong evidence in favour of the null hypothesis (i.e. that the mPFC is neither more or less activated in the False compared to True Belief condition).

#### 3.2.3 Lateralisation of TPJ activity in spontaneous ToM

Investigation of lateralisation revealed a significant interaction between location and agent (*F*(1,67) = 19.95, *p* < 0.001) meaning that the right TPJ was more active than the left TPJ when the agent believed the ball was present (A+) than when he believed the ball was absent (A-). Another significant interaction emerged between location, agent and participant (*F*(1,67) = 7.49, *p* = 0.008) meaning that the difference in activation between the False and True Belief conditions was stronger in the right TPJ than in the left TPJ. A post hoc paired t-test revealed significantly higher activity in the right TPJ as compared to the left TPJ in the False Belief condition (*t*(67) = 2.818, *p* < 0.05). A repeated-measure ANOVA with location and content of False Belief as factors revealed a main effect of location (*F*(1,67) = 7.94, *p* = 0.006) and a significant interaction between location and content (*F*(1,67) = 13.42, *p* < 0.001). Post hoc t-tests revealed significantly higher activation of the rTPJ, but not of the lTPJ, in the False Belief with positive content condition as compared to the False Belief with negative content condition (P-A+ versus P+A-; rTPJ, *t*(67) = 3.06, *p* < 0.01; lTPJ, *t*(67) = 0.60, *p* = 0.553). These results confirm the asymmetry of the right TPJ, but not of the left TPJ, in response to content in the False Belief condition.

#### 3.2.4 Cognitive decoding of brain activity

To evaluate the consistency of our findings with the average pattern of activation associated with ToM, we compared the maps obtained in the PxA analysis with the activation maps stored by Neurovault. The results are displayed in Figure 5 for both contrasts. The activation map for the False > True Belief contrast corresponded more strongly to the activation maps related to working memory (all structures: *r* = 0.23; cortex: *r* = 0.24; subcortex: *r* = 0.10) and visuospatial functions (all structures: *r* = 0.14; cortex: *r* = 0.15; subcortex: *r* = −0.01) than to ToM (all structures: *r* = −0.04; cortex: *r* = −0.06; subcortex: *r* = −0.07), especially in the more dorsal active areas, according to visual inspection.

**Figure 5.**
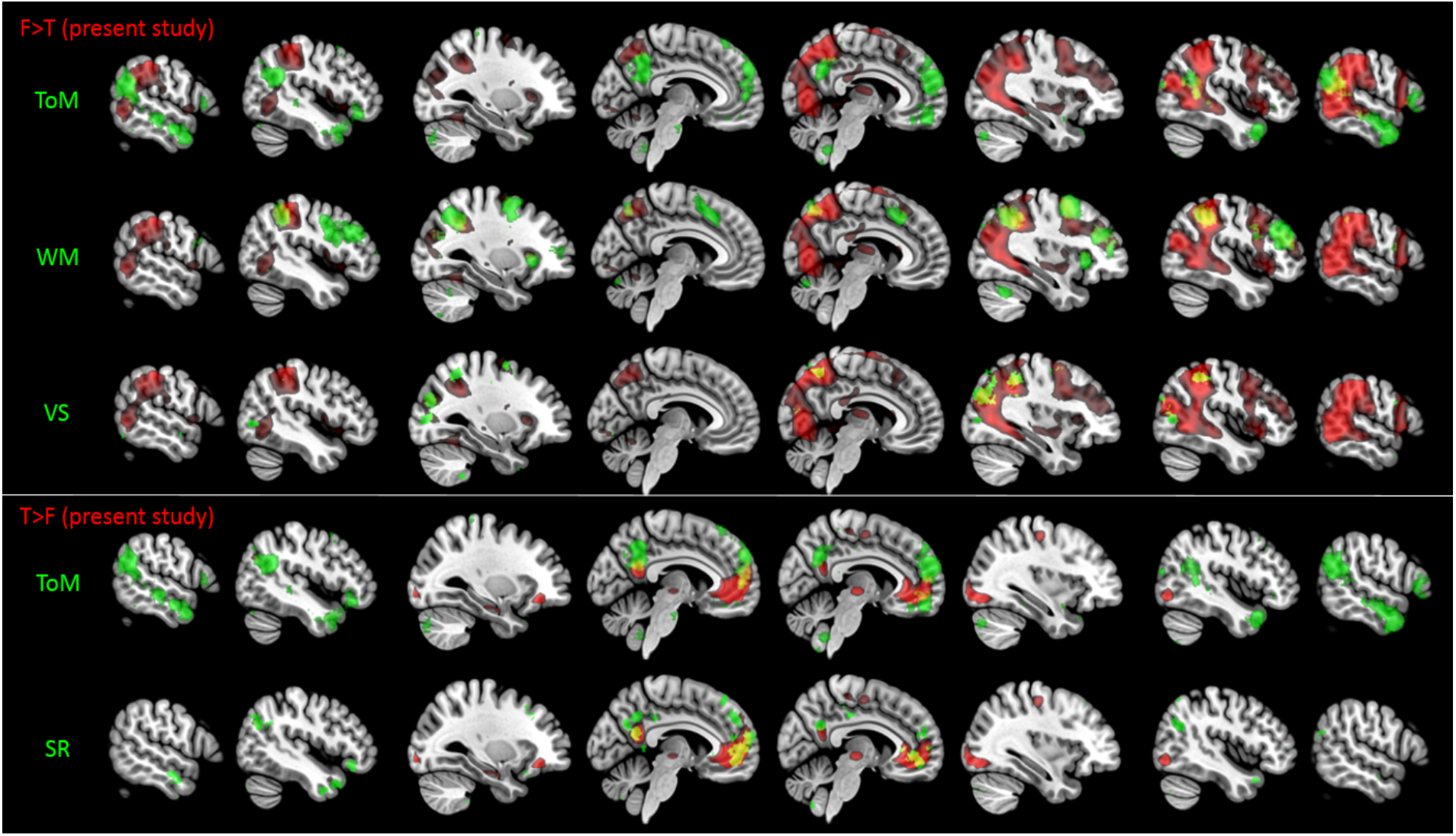
Overlap with Neurosynth maps. Figures were visualised with MRIcroGL software (https://www.mccauslandcenter.sc.edu/mricrogl/). Red colour corresponds to our activation map for the PxA analysis in all the figures. First three rows from the top correspond to the False > True Belief (F>T) condition, last two rows to the True > False (T>F) condition. Green always corresponds to a map of Neurosynth. Yellow corresponds to the overlap between our map and Neurosynth map. From the top to the bottom: overlap with Theory of Mind (ToM) map, working memory map (WM), visuo-spatial map (VS), ToM and self-referential map (SR).

#### 3.2.5. Brain-Behaviour Relation

To investigate the correlation between brain activity in the belief formation phase and behaviour in the outcome phase, we first added the RT ToM Index of each participant as a covariate to the models testing the False > True Belief contrasts, both for the PxA and ToM Index analysis. This yielded no significant effects. We then calculated the correlation between RT ToM Index and the neural ToM Index for the bilateral TPJ. Results are displayed in supplementary material (supplementary Figure 1). This revealed a borderline significant negative correlation in the rTPJ (*r* = −.206, 95% CI: [−.426 .038], *p* = 0.098) and in the lTPJ (*r* = −.237 [−.453 .005], *p* = 0.055) suggesting that a larger ToM index (i.e., increased mentalisation ability) moderately correlated with less right and left TPJ activity. However, after removing the outliers emerging from an outlier analysis, the correlations were no longer significant (rTPJ: *r* = −.119 [−.356 .133], *p* = 0.352; lTPJ: *r* = −.160 [−.391 .089], *p* = 0.206).

## 4. Discussion

This study sought to define the neural mechanisms of spontaneous ToM by pooling data from three studies in a mega-analytic fashion to better identify the brain areas involved in spontaneous ToM and to resolve some of the present controversies. Partly confirming our first hypothesis, spontaneous false belief processing during the belief formation phase activated a large cluster spanning the temporo-parietal cortex, and including the TPJ, the same region activated in explicit false belief processing. Nevertheless, no activation in the mPFC, a region involved in the explicit ToM, for the False > True Belief contrast was observed. ROI analysis confirmed these results, showing significant activation of bilateral TPJ and strong evidence against the involvement of the mPFC during the belief formation phase. Thus, the spontaneous ToM system partially overlaps with the explicit ToM system in terms of TPJ but not mPFC activation. Moreover, while our results showed that the rTPJ was active for both False Belief with positive and with negative content conditions, activation in the positive content condition was significantly higher than in the negative content condition, consistently with the hypothesis of rTPJ content-selectivity in spontaneous ToM. ROI analyses confirmed that both the right and left TPJ were activated by inconsistencies between the belief of the participant and the belief of the agent but with a dominance of the rTPJ. Lastly, the comparison with Neurosynth maps revealed that our maps of activation overlapped more with maps related to other functions (working memory and visuospatial for the False > True Belief contrast and self-referential for the True > False Belief contrast) than to ToM maps.

The main motivation for this study was to resolve prior discrepancy in fMRI studies of spontaneous ToM. Previous findings have been contradictory, especially regarding the role of the TPJ (Bardi et al., 2017; Kovács et al., 2014; Schneider et al., 2014) and consequently regarding whether the spontaneous and the explicit ToM networks overlap. With regards to the first main finding, in agreement with Kovács et al. (2014) and contrary to Schneider et al. (2014), our whole-brain analysis revealed a large cluster of active brain areas encompassing the TPJ in the False > True Belief contrast. The TPJ is a brain region consistently found to be active during explicit ToM and especially active in explicit false belief tasks (Döhnel et al., 2012; Saxe, 2010). Our finding provides evidence for an overlap between explicit and spontaneous ToM during the belief formation phase, suggesting that the TPJ is recruited both when participants are explicitly reasoning about others’ mental states and when they are spontaneously doing so. The subsequent ROI analysis confirmed that the rTPJ was activated by inconsistencies between the belief of the participant and the belief of the agent.

However, our results clearly indicate that the mPFC is not involved in false belief processing in a spontaneous ToM task, at least for the tracking of beliefs formation. This contrasts with substantial evidence that mPFC is consistently active in explicit ToM (Schurz et al., 2014; Van Overwalle, 2009). Until now, the absence of mPFC activation during belief formation in spontaneous ToM could have reflected a power issue that left the activation undetected, or a real lack of mPFC involvement. Our analysis provides enough statistical power to prove that the mPFC is actually not recruited during spontaneous mentalising. When considering the implications of these findings it is crucial, however, to take into account that we measured brain activation in the belief formation phase while most explicit ToM studies measure activation when participants report the beliefs. It might well be that the mPFC is related to reporting beliefs, i.e. during later stages of cognitive processing and that even in explicit ToM tasks the mPFC is not found to be active in the belief formation phase, as reported by Bardi and colleagues (2017). This argues against a dual process perspective.

The same explanation can account for the fact that our results contradict those by Schneider and colleagues (2014), who reported activation of the left STS and the precuneus, but not the TPJ. Indeed, their analysis was carried out on the outcome phase rather than the belief formation phase, as in the present study. Therefore, it is possible that the TPJ is a crucial region involved in belief processing, at least at the belief formation level.

The second question was whether we could replicate the content-selectivity of TPJ activation found by previous studies on smaller samples (Bardi et al., 2017; Kovàcs et al., 2014; Nijhof et al., 2018), who reported that the right TPJ was more active in the False Belief with positive content condition. Our data confirmed the content-selectivity of the right TPJ, although showing that the rTPJ was also active in the False Belief with negative content condition to a lesser degree. This representation limit of spontaneous ToM during the belief formation phase implies that when another person’s belief of a positive content attribution differs from our own belief, there is more activation of the spontaneous ToM system than when the other’s belief has a negative attribution. In other words, situations in which other people’s false beliefs are about the presence, and not the absence, of an object may be more relevant for an immediate prediction of others’ behaviour. Nevertheless, since the task required participants to press the button when the ball was present in the outcome phase, it is not possible to exclude that the bias for agent’s beliefs with positive content might reflect the task instructions.

A third central question was whether there is a functional dominance of the right TPJ over the left TPJ or if both sides are equally activated by the ToM task (in support of rTPJ dominance: Aichhorn et al., 2009; Döhnel et al., 2012; Liu et al., 2009; Saxe, 2010; in support of equal bilateral TPJ activation: Jenkins & Mitchell, 2010; Krall et al., 2015; Saxe and Kanwisher, 2003). Consistent with the main hypothesis, rTPJ relative to lTPJ activity was indeed statistically stronger although both regions were active during the False > True Belief condition. An additional ROI analysis also revealed a stronger False > True Belief effect in the right TPJ than in the left TPJ supporting a stronger right-lateralisation. These findings are in agreement with previous reports of rTPJ dominance in explicit ToM (Aichhorn et al., 2009; Döhnel et al., 2012 Liu et al., 2009; Saxe, 2010) and spontaneous ToM (Bardi et al., 2017; Hudson et al., [preprint, bioRxiv]; Kovács et al., 2014; Nijhof et al., 2018), although not excluding an involvement of the lTPJ. Yet the precise underlying functional mechanisms regarding this effect remain unknown.

Importantly, the TPJ was not the main source of brain activity and the main peaks of activation in the whole-brain analysis were located more dorsally and ventrally. The ventral peak corresponded to the middle temporal gyrus (MTG), a region implicated in ToM (Carrington & Bailey, 2009; Schurz et al., 2014) and involved in false belief processing (Rothmayr et al., 2011; Sommer et al., 2007; van Veluw & Chance, 2014). The dorsal peak corresponded to the supramarginal gyrus (SMG), or BA40, a region that is part of the inferior parietal lobule (IPL) (Igelström & Graziano, 2017) and is likewise activated in false belief tasks (see the meta-analysis by Schurz, Aichhorn, Martin & Perner, 2013). However, the SMG is also involved in other functions, such as spatial working memory, spatial attention and visuospatial processing (Silk, Bellgrove, Wrafter, Mattingley & Cunnington, 2010; Walter & Dassonville, 2008). Therefore, our findings, while confirming the activation of the TPJ in false belief reasoning, also extend this role to other parietal and temporal areas with peaks in the SMG and MTG, respectively. However, activation in frontal areas also emerged, namely, the right middle and inferior frontal gyri (MFG, IFG). These findings are in agreement with those reported in previous research finding MFG activity during both spontaneous (Naughtin et al., 2017) and explicit ToM (Rothmayr et al., 2011; Sommer et al., 2007). The IFG is another area involved in attentional mechanisms, especially attention switching (Hedge et al., 2015), thus activating when a discrepancy in the observed situation, such as a belief of another person that differs from our own, catches our attention.

To better integrate findings from the present analysis with the broader literature, the maps for the False> True Belief were compared to available maps collected by Neurosynth. This comparison revealed that activation obtained for the False > True Belief contrast corresponded most strongly to activation maps of visuospatial and working memory functions, especially in dorsal regions, and had only little resemblance with the typical activation pattern found in ToM tasks. The main dorsal peak of activation found in our analysis corresponded to the SMG, a region, as said, that is involved in spatial working memory and visuospatial processing (Silk et al., 2010; Walter & Dassonville, 2008). Since false belief processing requires individuals to keep in mind the belief of another person, working memory may be needed to correctly perform the task and thus regions involved in this function might be recruited when participants are spontaneously reasoning about others’ mental states. Moreover, the present task implemented in this study required visuospatial processing, since participants had to track the location of a ball. When participants have a certain knowledge on the position of the ball that is different from that of the agent (false belief conditions), visuospatial areas might be recruited more because of the effort provided by knowing where the object actually is and simultaneously spontaneously imagining the ball in the location where the agent believes it to be. The dorsal peak of activation found in this study might therefore be involved in remembering and tracking the different assumed location of the ball, but still be important to correctly carry out the ToM task in the false belief conditions. As such, visuospatial functions may be recycled to monitor the beliefs of others (Corbetta, Patel G & Shulman, 2008) and so the dorsal activation could reflect real ToM activity and not be an artefact of the task.

The activation of regions involved in visuospatial processing, working memory and attention shifting, such as the SMG and the IFG, might suggest that spontaneous ToM actually is simply a low-level general spatial processing function, based on submentalising domain-general mechanisms, that is activated when the observed situation presents a discrepancy between one’s own and anothers’ point of view, perspective or belief (see review by Heyes, 2014). This discrepancy would be processed in a spontaneous, fast and efficient way without the need to inhibit one’s own belief. Inhibition only comes into play when participants have to explicitly respond to a question during the task, thus requiring the involvement of the explicit ToM system, which includes the mPFC.

However, by overlapping the map for the False > True Belief with the maps of Neurosynth, we were able to compare our map with ToM maps obtained by many different studies that used different ToM tasks. Thus, it is important to bear in mind that those ToM tasks might activate different brain regions than the task used in the present study, which might explain why we found little overlap with the Neurosynth maps.

Lastly, the behavioural analysis confirmed that the agent’s belief had a biasing effect on participants’ responses, which were faster when the agent believed the ball was present than when the agent believed the ball was absent. The correlation between behaviour in the outcome phase and brain activity in the belief formation phase, however, was not significant after removing the outliers.

Crucially, the bigger sample of the present study, compared to previous work, increased the statistical power of fMRI analysis, allowing us to overcome the power issue that constitutes a strong limitation of previous findings. Thus, it is possible to confirm that the absence of mPFC activity during belief formation in spontaneous ToM, which was also reported in previous studies, is not due to poor statistical power but reflects a characteristic of the spontaneous ToM network. Hence, the increased power provided by the mega-analytic approach implemented in the present study constitutes its main strength and grants more confidence in the findings.

## 5. Limitations

Despite these intriguing findings, it is important to consider that the study did not include a functional explicit ToM localiser as comparison. Including a ToM localiser would be important in order to better elucidate how the pattern of activity in the temporo-parietal cortex found in the False > True Belief conditions overlapped with explicit ToM areas in a more direct way than with ToM maps in Neurovault and also to better define the ROIs, rather than using coordinates from a meta-analysis. Another limitation comes from the fact that spontaneous ToM studies require a focus on the belief formation phase, while explicit ToM studies mainly focus on the outcome phase. As such, comparing the explicit and spontaneous ToM systems is tricky due to the intrinsic nature of the tasks.

## 6. Conclusions

In conclusion, the present mega-analysis sought to characterise the network involved in spontaneous ToM and better understand whether spontaneous and explicit ToM rely on the same or distinct networks. The main analysis revealed a large cluster of activation related to spontaneous false belief processing in the right temporo-parietal cortex, encompassing the right TPJ and two main peaks in more dorsal and ventral locations, probably related to visuospatial and working memory functions that might be useful for carrying out the type of task used in this study or which might be inherent to ToM (see review by Corbetta et al., 2008). No mPFC activation emerged. The ROI analysis confirmed these findings. Thus, explicit and spontaneous ToM networks overlap regarding the role of the TPJ during the formation of beliefs, but not for the mPFC. Content-selectivity of the spontaneous ToM network and asymmetry in rTPJ versus lTPJ activation were confirmed, while no effects of functional connectivity emerged. Comparison with available maps suggested overlap with a variety of higher-order cognitive functions.

Overall, these findings suggest that the explicit and spontaneous ToM systems partially overlap, showing similar activation (TPJ) in the belief formation phase. However, spontaneous mentalising does not recruit the mPFC at the level of beliefs formation. In conclusion, the spontaneous ToM seems to be based on domain-general mechanisms, which provide a fast, efficient response to salient social stimuli, like discrepancies in belief or perspective, via low-level general spatial processing functions.

## Acknowledgements

Sara Boccadoro would like to thank the Erasmus+ Traineeship program for the travel bursary.

## Funding

The study by Bardi et al. was funded by Research Foundation - Flanders (FWO) Pegasus Fellowship to Lara Bardi and by grant 331323-Mirroring and ToM, Marie Curie Fellowship (Marie Curie Intra-European fellowship for career development) to LB. The study by Nijhof et al. was was supported by the Special Research Fund of Ghent University (project number BOF13/24J/083). The study by Hudson et al. was supported by a 2-4 year grant (01J05415) from the Special Research Fund (BOF) at Ghent University to SCM and MB.

## Conflict of interest

Declarations of interest: none.

## Supplementary material

### PPI analysis

The gPPI analyses examined brain-wise functional connectivity of 1) the a priori rTPJ ROI and 2) of the two main temporo-parietal peaks that emerged from the whole-brain analysis (rMTG and SMG) with the rest of the brain for the False > True Belief contrast. No significant findings emerged.

### PPI analysis discussion

Based on very limited evidence from functional connectivity analysis during a social emotion task (Burnett & Blakemore, 2009), we predicted functional connectivity between the TPJ and the anterior mPFC. However, the PPI analyses did not reveal any significant findings, thus failing to characterise the functional network of spontaneous false belief processing.

**Supplementary figure 1.**
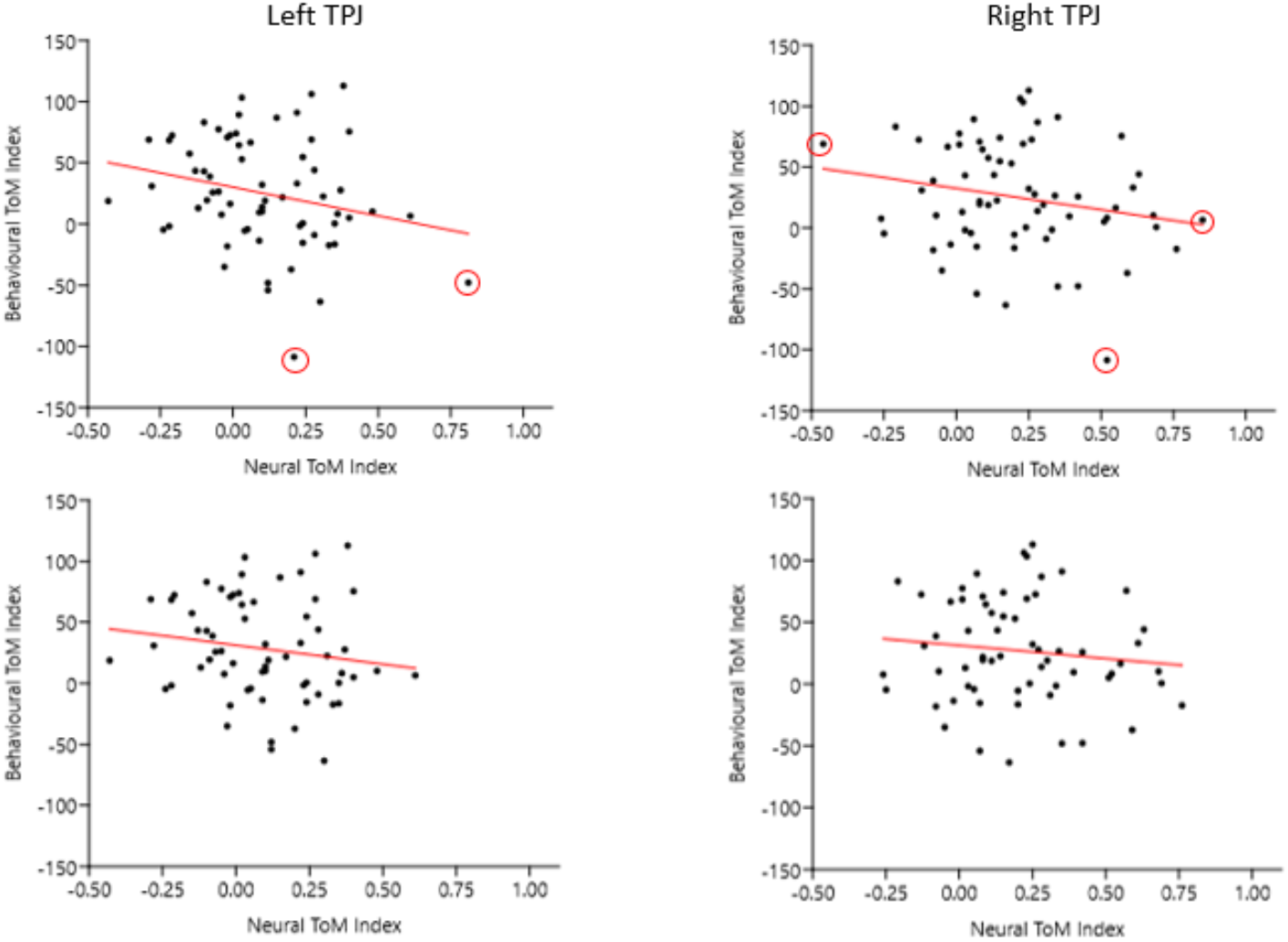
Correlation between behavioural and neural ToM Index. The scatter plots were created with PAST statistical software (https://folk.uio.no/ohammer/past/; Hammer, Harper & Ryan, 2001). The figure shows the correlation graphs of the left TPJ (on the left) and right TPJ (on the right) neural ToM Index with the behavioural ToM Index, with (on the top) and without (on the bottom) outliers. The outliers are circled in red.

